# Arterial Elastin Abundance, Rather Than Orthologue Origin, Modulates Medial Arterial Calcification in Matrix Gla Protein-Deficient Mice

**DOI:** 10.64898/2026.08.31.748131

**Authors:** Juliana Marulanda, Ophélie Gourgas, Abhinav Parashar, Robert P. Mecham, Elaine C. Davis, Marta Cerruti, Juergen Brinckmann, Monzur Murshed

**Author notes:** Corresponding author: Monzur Murshed, PhD, Address: Shriners Hospital for Children, 1003 Boulevard Décarie, Montreal, QC, H4A 0A9 Canada.

## Abstract

Calcific deposits in the arterial media have been associated with a number of metabolic and genetic disorders including diabetes, chronic kidney disease and generalized arterial calcification of infancy. While medial calcification and physiologic ‘hard’ tissue mineralization in the skeleton are both regulated by several common determinants, emerging data suggest that there might be fundamental differences in the mechanisms underlying these two processes.

**Objective:** We previously demonstrated that elastin haploinsufficiency delays medial calcification in MGP-deficient mice. Here, using mice in which a human *ELN* transgene rescues mouse elastin deficiency, we investigated whether the origin and abundance of arterial elastin differentially affect the initiation and progression of medial calcification.

**Approach and Results:** We pursued a transgenic approach to alter the arterial elastin scaffold in MGP-deficient mice. Our analyses of a humanized MGP-deficient model with 40% reduction of medial elastin content showed a complete absence of the early-stage vascular calcification. Additionally, we showed that mouse and human elastin orthologues affect vascular calcification in a comparable manner.

**Conclusion:** Arterial elastin abundance, rather than orthologue origin, modulates the initiation and progression of medial calcification in MGP-deficient mice. A further reduction in arterial elastin beyond that achieved by elastin haploinsufficiency profoundly delays mineral deposition and maturation, whereas restoration of elastin abundance through transgenic human ELN expression restores arterial calcification.

## Introduction

ECM mineralization is a physiologic process in the skeletal ‘hard’ tissues, but a pathologic condition, when occurs in the ‘soft’ tissues like the blood vessels. Both physiologic and pathologic mineralization require a suitable ECM that can act as a scaffold for the nucleating minerals. Additionally, the process is regulated by the extracellular levels of mineral ions such as calcium and inorganic phosphate and various mineralization inhibitors^1,2^.

Ectopic mineral deposition in vascular tissues is commonly known as vascular calcification and has been traditionally considered as a passive process, a corollary of aging or degenerative diseases such as advanced atherosclerosis, chronic kidney disease (CKD) and type II diabetes^3^. However, more recent publications consider vascular calcification as an active and regulated process that shares many common determinants with the physiologic mineralization process of bones and teeth^2^.

Depending on the anatomical location, vascular calcification in the blood vessels has been classified into two types: intimal and medial calcification. The former is typically associated with atherosclerosis, showing dispersed, punctuate or patchy mineral crystals associated with the necrotic core of the atheroma. On the contrary, in medial calcification, minerals are deposited within and around the medial elastic lamina, which can develop independent of atherosclerotic lesions^3,4^

In the current study, we investigated the epistatic effect of elastin, a key genetic regulator of ECM mineralization affecting the initiation and progression of medial calcification in MGP-null (*Mgp−/−*) mice^5^. We used complex genetic models to examine how genetic modifications involving the dosage of elastin (*Eln*) may change the severity of vascular calcification phenotype in *Mgp−/−* mice.

MGP is among the most potent mineralization inhibitors expressed mainly in the vascular and cartilaginous tissues. Mutations in the *MGP* gene in humans lead to Keutel syndrome, a rare disease hallmarked by abnormal cartilage and vascular calcification^6–8^. More recently, our laboratory reported dominant heterozygote variants of MGP (cys19phe or cys19tyr) causing spondyloepiphyseal dysplasia, MGP type in humans. While the affected individuals and the mouse model expressing the cys19phe variant manifest some overlapping skeletal pathologies as seen in Keutel syndrome, they do not show any vascular calcification^9^.

The *Mgp−/−* mice recapitulate most of the traits seen in patients with Keutel syndrome, but with a more severe vascular calcification phenotype^5^. The presence of calcific deposits in arterial tissues in these mice can be detected as early as 5 days after birth, and after that, the amount of deposited minerals increases progressively^5^. Our previous work has demonstrated that the early phases of medial calcification in *Mgp−/−* mice do not require any upregulation of chondrogenic/osteogenic genes in the aorta^10^.

We reported earlier that the reduction of the elastin gene dosage in *Mgp−/−;Eln+/−* mice caused a delay of mineral deposition in the arterial media^10^. However, the progressive increase of deposited minerals continued and eventually these mice succumbed to this phenotype. It remains unknown whether a further reduction in arterial elastin would more profoundly delay vascular calcification and whether human and mouse elastin orthologues have comparable capacities to support mineral deposition.

In the current study, we used a genetic approach to show that an approximately 40% reduction in medial elastin content in an MGP-deficient compound-mutant model prevents detectable early arterial calcification and profoundly delays its subsequent progression. Further, we provide evidence that the mineral scaffolding properties of mouse and human orthologues of elastin are conserved and comparable. This study contributes to our understanding of the mechanisms regulating medial vascular calcification.

## Material and Methods

### Mouse models

The generation of *Eln+/−(BAChELN)* mice (hereinafter referred to as *Eln+/−; ELN+/+*) has already been described^5,11^. All the animal experiments were performed according to the animal use protocol number 7132 approved by the Animal Care Committee of McGill University.

### Micro-CT and mineral quantification

Micro-CT scanning of mouse aortas were performed with a SkyScan model 1272 instrument (SkyScan) set at a resolution of 8.0μm and 0.25-mm Al filter. Image acquisition was performed at 40Kv and 250µA with a 0.4-degree rotation between frames. Adaptive thresholding was applied to bone and aorta separately. Micro-CT image processing and analysis was performed with version 2.2f of the manufacturer’s software (SkyScan). The 3D-reconstructions were done using the CtAn and CtVol software (SkyScan). Mineral quantification was calculated using CtAn software (SkyScan), considering the BV/mm^3^ generated in the basic values.

### Histology and tissue imaging

Mouse thoracic cages were fixed overnight in 10% formalin, embedded in methyl methacrylate, sectioned (7μm), and stained by von Kossa and van Gieson (VKVG) to visualize calcified tissues and collagen, Hart’s staining to visualize the elastic lamina; or Alcian blue and van Gieson (ABVG) to detect proteoglycans in the cartilaginous matrix. Images were taken using a light microscope (DM200; Leica Microsystems) with 20X (numerical aperture of 0.40) and 40X (numerical aperture of 0.65) objectives. All histological images were captured using a digital camera (DP72; Olympus Canada Inc), acquired with DP2-BSW software (XV3.0; Olympus Canada Inc) and processed using PhotoShop software (Adobe).

### Serum biochemistry

Serum Ca^2+^ and P_i_ levels were measured using commercially available kits (Sekisui Diagnostics P.E.I. Inc, Charlottetown, Canada).

### Tissue biochemistry

For collagen and elastin analysis, specimens were cut into small pieces and digested with high-purity bacterial collagenase (C0773; Sigma, Germany; 1U/mL, 37°C, 12 hours). After centrifugation, the soluble fraction containing collagen was subjected to hydrolysis and amino acid analysis. The residual fraction was extracted by hot alkali (0.1 N NaOH, 95°C, 45 minutes). After centrifugation the supernatant containing non-collagenous/non-elastin proteins (NENCs) and the insoluble residue containing insoluble elastin were subjected to hydrolysis. Hydrolysis was performed in 6 N HCl at 110°C for 24 hours. Dried hydrolyzates were redissolved in sodium citrate loading buffer (pH 2.2) and amino acid analysis was performed by ion exchange chromatography with postcolumn derivatization with ninhydrin (Biochrom 30; Biochrom, United Kingdom). The amino acid profile of the supernatant after collagenase digestion showed a profile typical for collagen, whereas the supernatants after hot alkali treatment were free of hydroxyproline. The amino acid profile of the insoluble fraction after hot alkali extraction was typical for elastin. The content of collagen, elastin, and of non-elastin/non-collagenous protein was expressed per total protein (µg/µg).

### Staining of thoracic aortae

Skeletal tissues from adult mice were fixed overnight in 95% ethanol, stained in 0.015% Alcian Blue dye (Sigma-Aldrich) in a 1:4 solution of glacial acetic acid and absolute ethanol for 24 hours. Tissues were then treated with 2% potassium hydroxide for another 24 hours (or until the soft tissues were dissolved) and then stained by 0.005% Alizarin Red (Sigma-Aldrich) in a 1% potassium hydroxide solution.

### Raman spectroscopy

Aorta sections were analyzed by Raman spectroscopy using a Bruker Senterra confocal Raman spectrophotometer equipped with a 785nm diode laser coupled with an Olympus optical microscope, using a 40x objective. Spectra were collected at 100 mW laser power and spectral resolution of 3.5 cm^-1^, between 400 and 1800 cm^-1^, with an integration time of 40 seconds and 2 co-additions. In total, 10 points per sample were analyzed. The data were analyzed using OPUS software (OPUS 7.0.0, Bruker, Karlsruhe, Germany).

### Scanning electron microscopy (SEM) and energy-dispersive X-ray spectroscopy (EDS)

Aorta sections were characterized using a combination of SEM and EDS. Samples were secured to an aluminum sample holder with carbon tape, and then coated with about 8 nm carbon (carbon sputter coater, EMS150R ES, Electron Microscopy Sciences (EMS)). The samples were then imaged using an Inspect-50 field emission SEM (FEI, Japan), at 10 kV operating voltage under high vacuum. EDS spectra were obtained in the regions of interest using an EDX spectrometer (EDX, Thermo Scientific, USA).

### Statistical analysis

All results are shown as means ± standard deviations. Statistical analyses were performed by Student’s *t* test or analysis of variance followed by Tukey’s multiple-comparison test and Dunnett’s multiple comparison test using GraphPad Prism software. Single asterisk indicates *p* <0.05, two asterisks indicate *p* <0.01, three asterisks indicate *p* <0.001 and four asterisks indicate *p* <0.0001.

## Results

### Comparative analyses of arterial elastin content in compound models expressing mouse and human elastin orthologues

In order to examine the effects of elastin orthologues and content on the initiation and progression of medial calcification, we used several genetic models expressing mouse and/or human elastin at various dosages. We first analyzed the precursor mouse strains which include 1) WT mice, carrying both copies of mouse elastin gene (*Eln*), 2) The haploinsufficient *Eln+/−* mice carrying only one allele of *Eln*, 3) The *Eln+/−;ELN+/+* (*Eln+/−, BAChELN)* mice, carrying one allele of *Eln* and two alleles of human elastin (*ELN*) transgene, and 4) The *Eln−/−;ELN+/+* mice, containing two alleles of *ELN* only. PCR analyses revealed that our breeding program resulted in all the above genotypes (**Figure 1A)**.

**Figure 1.**
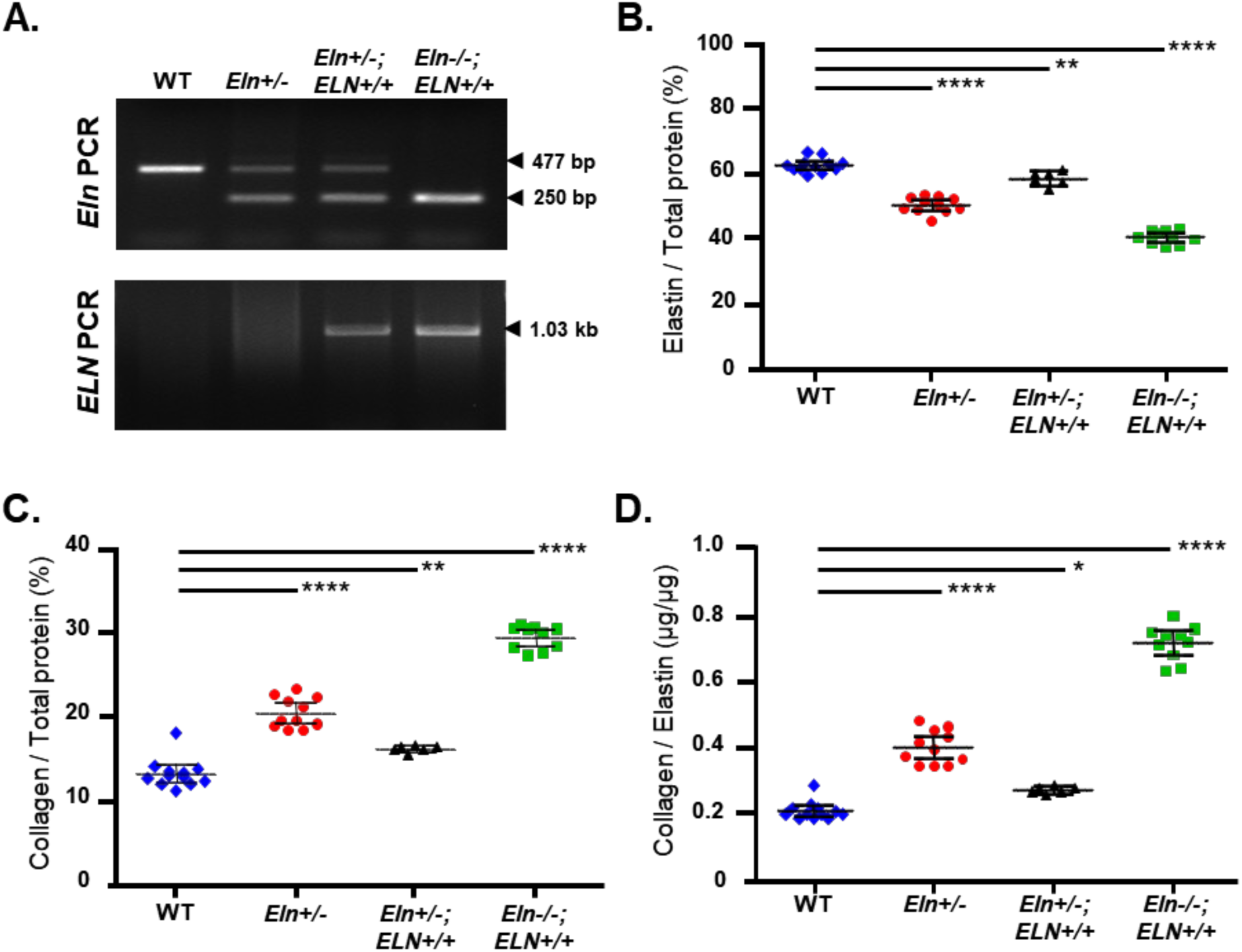
Arterial elastin content in mice expressing mouse and human elastin orthologues. **A.** PCR performed on the genomic DNA confirms the genotype of WT, *Eln*+/−, *Eln*+/−;*ELN*+/+ and *Eln*−/−;*ELN*+/+ mice. Mouse elastin (*Eln*) WT band is 477 bp, “knockout” band is 250 bp (top panel), and the human elastin transgene band is 1.03 kbp (lower panel). Homozygosity of the human transgene was determined by the survival of the *Eln*−/−;*ELN*+/+ mice, as all *Eln*−/− and *Eln*−/−;*ELN*+/− die perinatally. ^16 **B.** Elastin-to-total protein ratio in the aorta was decreased for all genotypes compared with WT mice, with the *Eln*−/−;*ELN*+/+ mice showing the lowest levels (n=6–12). **C.** Collagen-to-total protein ratio was elevated in all genotypes compared with WT, compensating for the loss of elastin. **D.** Collagen-to-elastin content (µg/µg) was the highest in *Eln*−/−;*ELN*+/+ aorta in comparison with WT. Statistical test: one-way ANOVA. n.s., not significant.

A comparison of the amino acid content of mouse and human elastin showed that the number of glycine and valine residues differs substantially between the orthologues (Supplemental material **Table SI** Supplemental table SI<u>.docx</u>. As expected, the relative numbers of these amino acids/1000 residues are comparable in WT and *Eln+/−* aorta. The amino acid composition and numbers in *Eln−/−;ELN+/+* aorta were comparable to the published human elastin data^11^. Further, the number of each amino acid residue in *Eln+/−;ELN+/+* aorta largely matched to its combined number in mouse and human elastin in 1: 1 ratio (Supplemental material **Table SI** Supplemental table SI<u>.docx</u>. The analysis of the amino acid composition showed that the breeding program resulted in the desired genotypes and the orthologue combinations.

We detected a decrease of elastin to total protein ratio for all genotypes compared to WT. The loss of elastin was the highest for *Eln−/−;ELN+/+* aorta and the lowest for *Eln+/−;ELN+/+* aorta (**Figure 1B**). The collagen to total protein ratio was elevated in all genotypes compared to WT. The highest and lowest ratios were found in the *Eln−/−;ELN+/+* and *Eln+/−;ELN+/+* aortas, respectively (**Figure 1C**). Based on these data, we calculated the collagen/elastin ratio in the arteries of various genotypes which was highest in the *Eln−/−;ELN+/+* mice (**Figure 1D**).

### Arterial mineral deposition is influenced by elastin abundance rather than orthologue combination

The milder vascular calcification trait in Keutel syndrome patients in comparison to the genetic model prompted us to investigate whether mouse and human elastin orthologues can promote medial calcification in a comparable manner. Earlier, we demonstrated that *Mgp−/−;Eln+/−* mice with a single allele of *Eln* could delay the mineralization in the arterial walls^10^. Based on the data presented in **Figure 1B**, elastin content in this model is expected to be approximately 25% lower than normal. In the current study, we examined whether the expression of the human orthologue in the arteries of *Eln+/−* mice, compensating for the loss of the endogenous protein can accelerate vascular calcification.

We bred the *Eln+/−;ELN+/+* mice with *Mgp+/−* mice to eventually generate *Mgp−/−;Eln+/−; ELN+/+* mice. The thoracic skeleton with the descending aortae from these and control *Mgp−/−* and *Mgp−/−;Eln+/−* mice were scanned and analyzed by micro-CT at 3 weeks of age (**Figure 2A**). As reported by us previously, we observed a reduction of the deposited minerals in the *Mgp−/−; Eln+/−* aorta in comparison to that of *Mgp−/−* aorta^10^. Interestingly, we observed more mineral deposition in the *Mgp−/−;Eln+/−;ELN+/+* aorta in comparison to that of *Mgp−/−;Eln+/−* aorta. In fact, the severity of the mineral deposition in the *Mgp−/−;Eln+/−;ELN+/+* and *Mgp−/−* aortae were visually indistinguishable. A quantitative analysis performed on the scanned images confirmed our qualitative observation (**Figure 2B**). We also observed a comparable mineralization of the elastic lamina by VKVG staining performed on histological sections of the aorta samples collected from the *Mgp−/−*, *Mgp−/−;Eln+/−* and *Mgp−/−;Eln+/−;ELN+/+* mice (**Figure 2C**).

**Figure 2.**
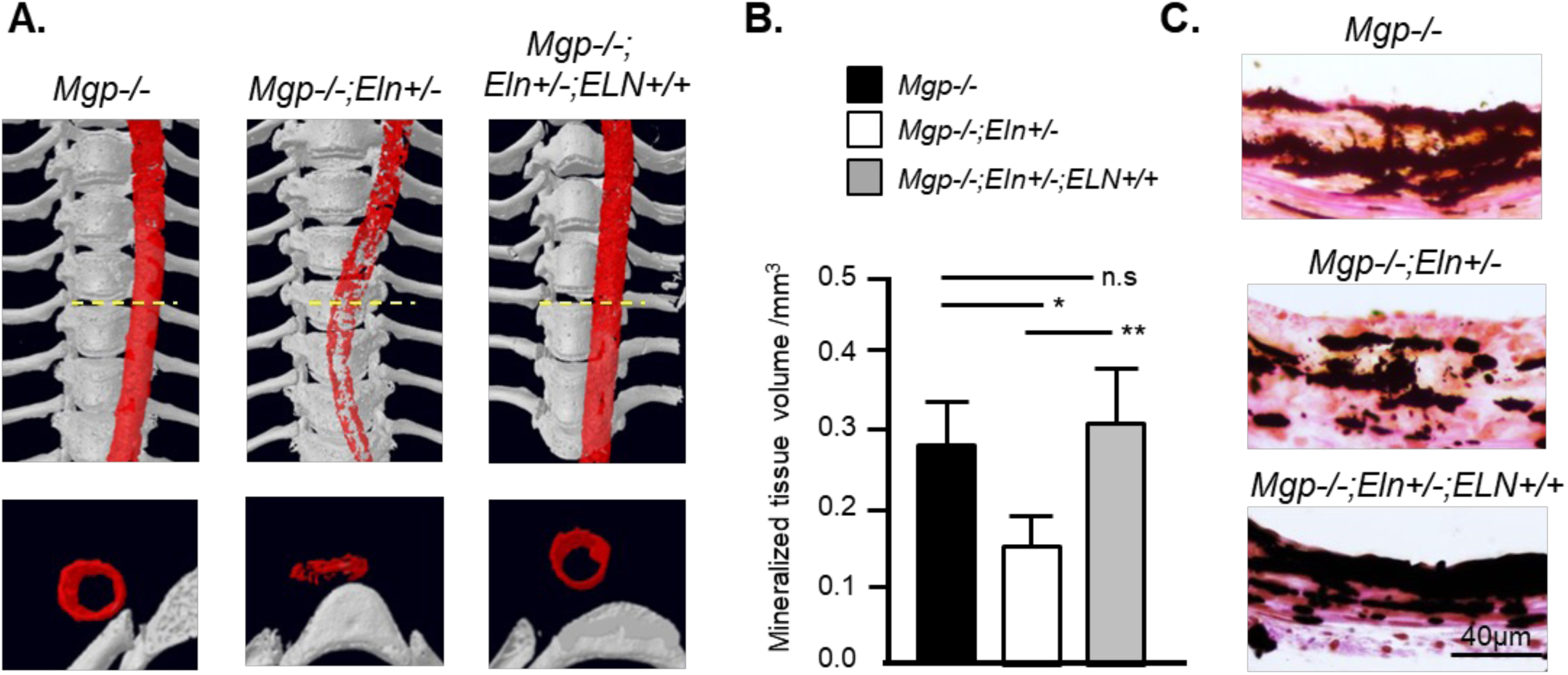
Vascular calcification is affected by the elastin content, but not by orthologue combination. **A.** 3D reconstructions of micro-CT scans showing severe thoracic aorta calcification in 3-week-old *Mgp*−/− mice. While *Mgp*−/−;*Eln*+/− mice showed a lower amount of minerals, age-matched *Mgp*−/−;*Eln*+/−;*ELN*+/+ mice producing a higher amount of elastin than the *Mgp*−/−;*Eln*+/− mice showed mineral deposition comparable to that of *Mgp*−/− mice (top panel). The lower panel shows the cross-sectional view at the yellow line. The aortic minerals were colored artificially in red. **B and C.** Aortic mineral quantification by micro-CT and VKVG staining of histological sections of the aorta confirms the data presented above (n=3). Statistical test: one-way ANOVA. n.s., not significant.

### Absence of detectable early vascular calcification in Mgp−/−;Eln−/−;ELN+/+ mice

The comparable abilities of mouse and human elastin orthologues to affect medial calcification established the humanized model (*Eln−/−;ELN+/+* mice) developed by Hirano *et al.*^11^ as a valid tool for further mechanistic studies on vascular calcification. Considering that the arterial elastin content is the lowest in these mice in comparison to the rest of the genetic models used by us, we decided to examine how this would affect the severity of medial calcification. The homozygous *Mgp*-null mutations were introduced to the humanized model to generate the *Mgp−/−;Eln−/−; ELN+/+* mice.

Remarkably, the *Mgp−/−;Eln−/−;ELN+/+* mice survive at least up to 10 months, the experimental end point of our survival study (**Figure 3A**) and beyond that period. Alizarin red and Alcian blue staining of the thoracic aorta revealed a complete absence of calcification in these mice at 3 weeks of age, while the aorta was heavily calcified in the age-matched *Mgp−/−* mice (**Figure 3B**). VKVG staining of thoracic aorta sections confirmed the absence of mineral deposition in the *Mgp−/−;Eln−/−;ELN+/+* mice. Additionally, the typical widening of the arterial wall seen in *Mgp−/−* aortas was also absent in the *Mgp−/−;Eln−/−;ELN+/+* aortas. However, we observed an increase of the collagen staining adjacent to the elastic laminae in the latter genotype. Hart’s staining showed that the elastic laminae were thinner, but more in number in comparison to the WT mice. Additionally, we also observed an increased cell number in the *Mgp−/−;Eln−/−;ELN+/+* mice (**Figure 3C**). As expected, serum P_i_ and Ca^2+^ levels remained unchanged (**Figure 3D**).

**Figure 3.**
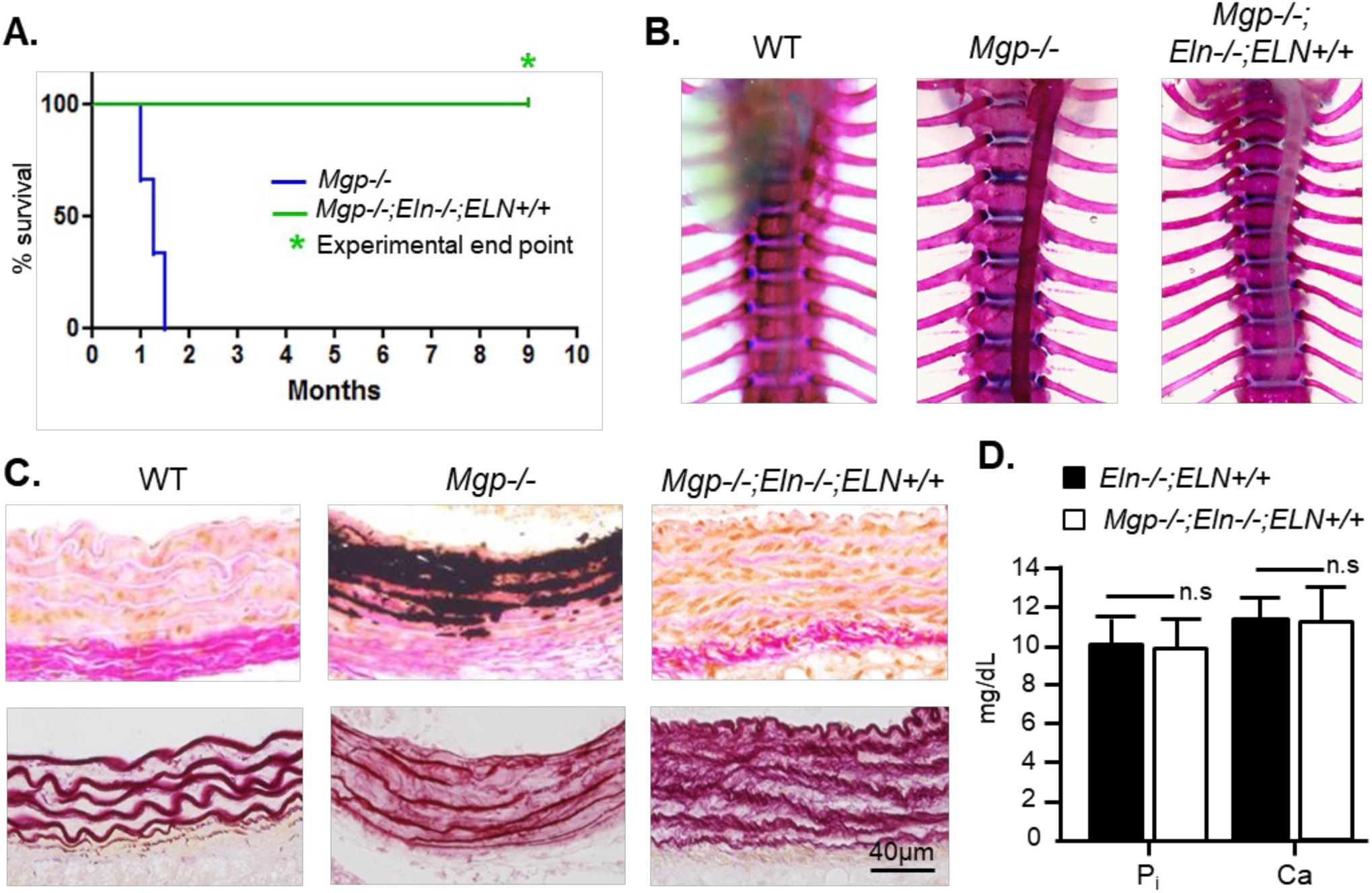
Absence of detectable early vascular calcification in *Mgp−/−;Eln−/−;ELN+/+* mice. **A.** Kaplan–Meier survival curve showing the survival of the *Mgp*−/−;*Eln*−/−;*ELN*+/+ mice until the experimental endpoint (10 months) (n=6 per genotype). **B.** Alizarin red and Alcian blue staining of the thoracic aorta shows severe vascular calcification in *Mgp*−/− mice, but remarkably, no vascular calcification was detected in the *Mgp*−/−;*Eln*−/−;*ELN*+/+ mice. **C.** Histological analysis of undecalcified plastic sections stained with VKVG (top panel) showing medial calcification in 3-week-old *Mgp*−/− thoracic aortas, but a complete absence of minerals in age-matched *Mgp*−/−;*Eln*−/−;*ELN*+/+ aortas. Note the augmented VSMC number (stained orange) and collagen staining (pink) in this latter model. Hart’s elastin staining of consecutive sections (lower panel) shows the loss of the wavy architecture of the elastic lamina in *Mgp*−/− aortas and an overall increased layering and decreased thickness of the elastic lamina in *Mgp*−/−;*Eln*−/−;*ELN*+/+ mice. **D.** Comparable serum P_i and Ca²⁺ levels in aged *Eln*−/−;*ELN*+/+ and *Mgp*−/−;*Eln*−/−;*ELN*+/+ littermates (n=5). Statistical test: Student’s *t* test. n.s., not significant.

We next examined the overall status of vascular calcification in these mice at 1 year of age. 3D reconstruction of micro-CT scans of the thoracic region of these aged mice revealed a mild and punctate mineralization throughout the thoracic aorta and in the coronary and intercostal arteries. In comparison, all these arteries are already heavily calcified by 5 weeks of age in *Mgp−/−* mice (**Figure 4A**). These findings were further confirmed by VKVG staining of the thoracic aorta sections using 1-year-old *Eln−/−;ELN+/+* aorta sections as controls. Hart’s staining of the elastic laminae revealed elastic fiber fragmentation and medial widening in the calcified *Mgp−/−;Eln−/−;ELN+/+* aortas (**Figure 4B**).

**Figure 4.**
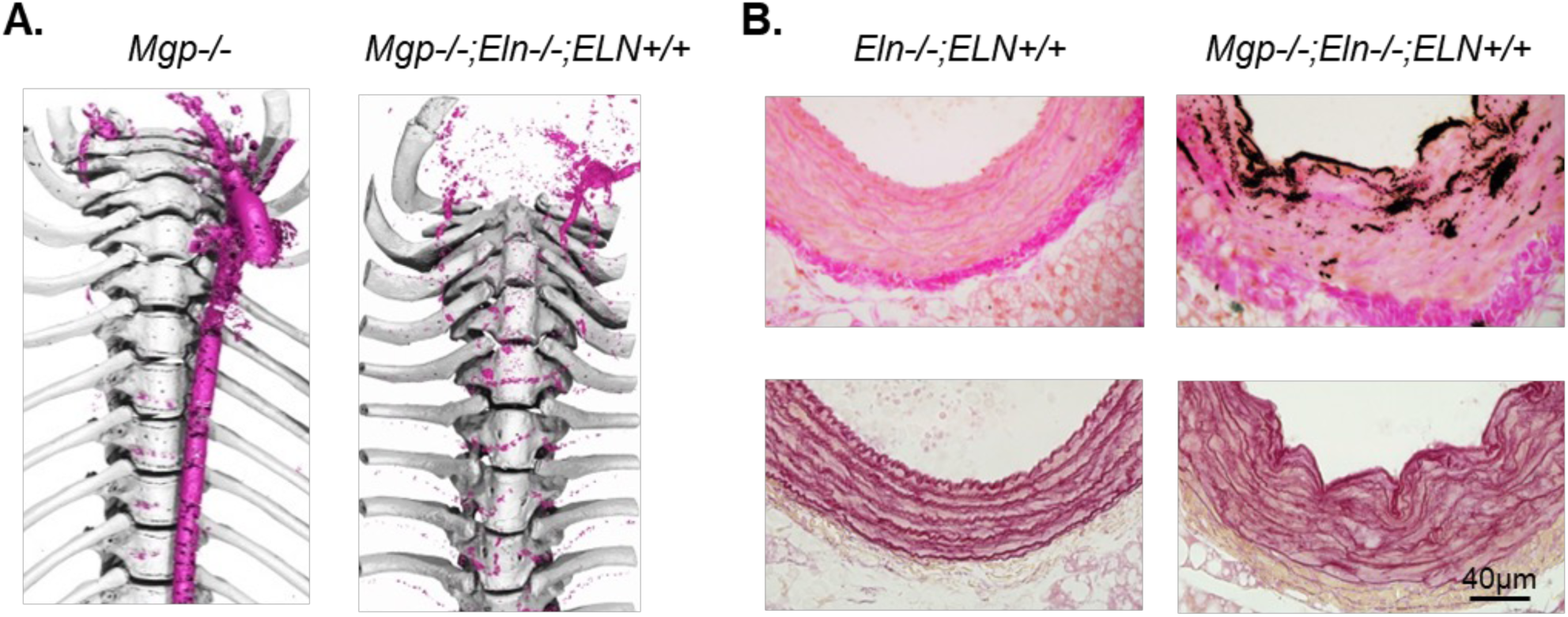
Aged *Mgp*−/−;*Eln*−/−;*ELN*+/+ mice develop mild vascular calcification. **A.** 3D reconstructions of micro-CT scans of the ribcages of *Mgp*−/− mice at 5 weeks of age and *Mgp*−/−;*Eln*−/−;*ELN*+/+ mice at 1 year of age comparing the extent of aortic mineral accumulation. Note the severity of vascular calcification in the *Mgp*−/− mice and the minor ectopic calcification in the latter model, where the coronary and intercostal arteries appear to be most affected. **B.** Histological sections of 12-month-old *Eln*−/−;*ELN*+/+ control and *Mgp*−/−;*Eln*−/−;*ELN*+/+ thoracic aortas stained with VKVG showing a mild deposition of minerals along the internal elastic laminae in the latter genotype (top panels). The lower panels show consecutive sections stained with Hart’s stain showing the overall reduction of elastic laminae in both models and the change in tissue architecture.

### Mineral characterization of the aortas from aged Mgp−/−;Eln−/−;ELN+/+ mice

We next used Raman spectroscopy in combination with SEM and EDS to characterize the mineral deposits present in the aged *Mgp−/−;Eln−/−;ELN+/+* mouse arteries. While Raman spectroscopy shows only the presence of CHA (◻_1_ phosphate band between 959 and 962 cm^-1^) in *Mgp−/−* arteries (**Figure 5A**), calcified *Mgp−/−;Eln−/−;ELN+/+* aortas show other mineral phases in addition to CHA (see the several υ_1_ phosphate vibration bands between 942 and 987 cm^-1^ on **Figure 5A** and **Table SII** Supplemental table SII<u>.docx</u>. Although Raman spectroscopy did not reveal precursor phases in *Mgp−/−* samples, using a combination of several microscopic and spectroscopic techniques we previously showed that the minerals are a mixture of precursor and apatite phases^12^. The difference in Raman sensitivity for detecting the precursor phases between the two mouse models suggests that the amount of those phases is likely much less prevalent in *Mgp−/−* than in *Mgp−/−;Eln−/−;ELN+/+* mice.

**Figure 5.**
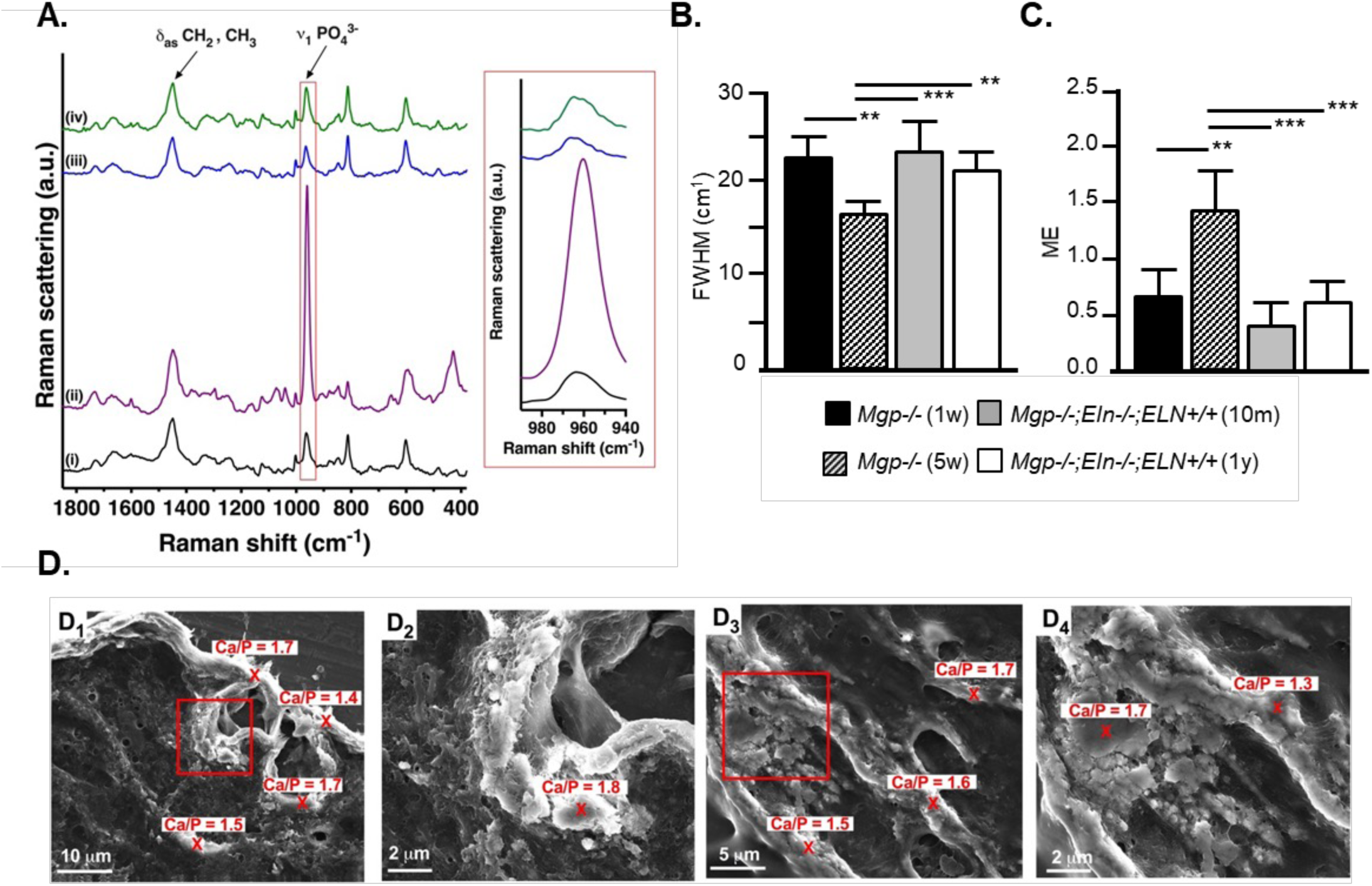
Mineral characterization of the aortas from aged *Mgp*−/−;*Eln*−/−;*ELN*+/+ mice. **A.** Raman spectra of calcified aortas from (i) 1– and (ii) 5-week-old *Mgp*−/− mice, and from (iii) 10-month-old and (iv) 1-year-old *Mgp*−/−;*Eln*−/−;*ELN*+/+ mice (n=3). **B.** FWHM values measured on the Raman ν₁ Pi peak. **C.** ME based on the Raman spectra. For each sample, 10 spectra were collected at different points. **D.** SEM images of calcified aortas from (D1 and D2) 10-month-old and (D3 and D4) 1-year-old *Mgp*−/−;*Eln*−/−;*ELN*+/+ mice. D2 and D4 are the insets in D1 and D3, respectively. The Ca/P ratios indicated on the SEM images are based on the elemental atomic percentages obtained by EDS spectroscopy. Statistical test: one-way ANOVA followed by Tukey’s test correction.

Regarding the crystallinity of the minerals, significantly higher FWHM values are found in *Mgp−/−;Eln−/−;ELN+/+* arteries compared to those found in 5-week-old *Mgp−/−* arteries while there is no significant difference with the FWHMs found in 1-week-old *Mgp−/−* arteries (**Figure 5B**). This suggests that the crystallinity of the minerals in the *Mgp−/−;Eln−/−;ELN+/+* mice is similar to the one of *Mgp−/−* mice at 1 week. The ME values in *Mgp−/−;Eln−/−;ELN+/+* mice are comparable to the values found in 1-week-old *Mgp−/−* mice and are significantly lower than the ones found in samples from 5-week-old *Mgp−/−* mice (**Figure 5C**).

SEM images of calcified aortas from 10-month-old and 1-year-old *Mgp−/−;Eln−/−;ELN+/+* mice show minerals in the form of compact material composed of smooth platelets, similar to the minerals present in *Mgp−/−* mice^12^ (**Figure 5D**). Ca/P ratios determined by EDS vary from 1.3 to 1.8, thus confirming that *Mgp−/−;Eln−/−;ELN+/+* samples contain several calcium phosphate phases, such as ACP, OCP, HA, and CHA^13^ (**Figure 5D**).

## Discussion

The similar pathological conditions associated with *ELN* mutations in humans and the genetically modified animals suggest that the functional properties of this protein are evolutionarily conserved^14^. This notion is further supported by the general pattern in the structure showing alternating hydrophobic and cross-linking domains in various elastin orthologues. Interestingly, although there is functional and structural conservation of elastin across the species, mouse and human elastin orthologues show only 64.1% identity at the amino acid level. This is much lower than the average 78.5% amino acid sequence identity between mouse and human proteins^14^. Despite this relatively low level of sequence identity, we found that the medial calcification in MGP-deficient mice, which were also heterozygotes for the mouse *Eln* gene and homozygotes for the human *ELN* transgene (*Mgp−/−;Eln+/−;ELN+/+* mice), mineralize comparably to that of *Mgp−/−* mice that carry two functional alleles of the mouse *Eln* gene. This result suggests a similar mineralization promoting capacity of the human elastin to that of mouse elastin. In the absence of any significant effect of orthologue types on the amount of deposited minerals, our data identify reduced elastin content as the major cause for the markedly reduced medial calcification in *Mgp−/−;Eln−/−;ELN+/+* mice.

Our published data suggest that MGP has a critical role to protect the arterial elastic laminae from the P_i_-induced mineral deposition^15^. This anti-mineralization property of MGP appear to be less critical in humans as the vascular calcification trait in Keutel syndrome patients is not as severe as in mice^7,8^. We show here that this milder trait may not be attributed to the sequence differences between the mouse and human elastin orthologues. At this point, the cause of a less severe vascular calcification phenotype in Keutel syndrome patients remains unknown. It is worth noting here that the basal P_i_ level in human blood is around 3.5 mg/dL as opposed to approximately 8 mg/dL in mice^16,17^. It is possible that the lower amount of circulating P_i_ in humans prevents the massive buildup of minerals in the human arteries lacking MGP. However, the involvement of additional regulatory mechanisms to prevent vascular calcification in humans cannot be ruled out.

Raman spectroscopy shows that mineral deposition in aged *Mgp−/−;Eln−/−;ELN+/+* mice is much less severe than in *Mgp−/−* mice at 5 weeks of age, when the arteries are severely calcified. Indeed, both the crystallinity and the amount of minerals found in *Mgp−/−;Eln−/−;ELN+/+* mice after 10 months are similar to those of the minerals in 1-week-old *Mgp−/−* mice. Both Raman and SEM/EDS data show that *Mgp−/−;Eln−/−;ELN+/+* samples contain a significant amount of precursor phases, such as ACP and OCP, in addition to apatite phases. These results confirm that although the nature of the minerals is generally similar, a 40% reduction of medial content of elastin in *Mgp−/−* mice significantly delays the initiation and progression of mineral deposition and slows down the maturation of the deposited minerals.

The presence of mineralization inhibitors PP_i_ and MGP protects the vascular tissues from spontaneous calcification. Interestingly, the presence of MGP was not sufficient to prevent vascular calcification caused by the reduced PP_i_ levels in GACI^18^. Similarly, in MGP deficiency, elastic lamina calcification occurs although the PP_i_ transport and synthesis mechanisms in the arterial tissues are both functional^10^. While the ectopic ECM mineralization caused by the absence of either of these inhibitors can be rescued by the reduction of the systemic P_i_ levels^15,19^, it remains to be seen whether they prevent the mineralization of different scaffolds in the arterial walls. Of note, despite an increase of the total collagen in the humanized arteries lacking MGP in *Mgp−/−; Eln−/−;ELN+/+* mice, no early calcification was associated with the collagen scaffold. It is possible that while MGP prevents the calcification of the elastic laminae, the anti-mineralization function of PP_i_ is restricted to the collagen fibers present adjacent to the elastic laminae. Further work will be needed to examine the complementary role of MGP and PP_i_ in the prevention of vascular calcification.

Although vascular calcification in *Mgp−/−* mice does not require the osteogenic or chondrogenic transdifferentiation of the VSMCs^10^, it does rely on several common determinants of ECM mineralization. Indeed, the basic tenet of skeletal tissue mineralization identifying the mineral ions, mineral scaffolding ECM and the absence of mineralization inhibitor as the key determinants is also valid for vascular calcification in MGP-deficient mice. Like bone mineralization, the vascular calcification in MGP-deficiency is P_i_-dependent and for its initiation, an elastin-rich extracellular scaffold is required and a mineralization inhibitor, MGP, must be removed or functionally inactivated^2,10,20^.

Our findings extend our previous observation that elastin haploinsufficiency delays medial calcification in MGP-deficient mice. Using a humanized elastin model, we show that a further reduction in arterial elastin profoundly delays the initiation and progression of mineral deposition and slows mineral maturation. Conversely, restoration of arterial elastin content through transgenic expression of human elastin restores the calcification-permissive phenotype. Thus, arterial elastin abundance has a greater influence than orthologue origin on MGP-deficiency-associated medial calcification. These findings identify the abundance and organization of the elastin-rich arterial scaffold as important determinants of susceptibility to medial calcification.

## Sources of Funding

The core facility for skeletal phenotyping was supported by Le Réseau de Recherche en Santé Buccodentaire et Osseuse (RSBO). This work was supported by following operating grants: CIHR 123310 to M.M and NIH HL53325 and HL105314 to RM.

## Disclosures

None. Authors have no competing interests to declare.

## Non-standard Abbreviations and Acronyms

ABVG: Alcian blue – Van Gieson
ACP: Amorphous calcium phosphate
ALPL: Alkaline phosphatase
BAC: Bacterial artificial chromosome
CHA: Carbonated hydroxyapatite
CKD: Chronic kidney disease
ECM: Extracellular matrix
EDS: Energy-dispersive X-ray spectroscopy
ELN: Elastin
FWHM: Full width at half maximum
GACI: Generalized arterial calcification of Infancy
HA: Hydroxyapatite
ME: Mineralization extent
MGP: Matrix Gla protein
Micro-CT: Micro-computed tomography
MVs: Matrix vesicles
NEXAF: Near-edge X-ray absorption fine structure spectroscopy
OCP: Octacalcium phosphate
OPN: Osteopontin
PHEX: Phosphate-regulating gene with homologies to endopeptidases on the X chromosome
PP_i_: Pyrophosphate
SEM: Scanning electron microscopy
VKVG: von Kossa – Van Gieson
VSMCs: Vascular smooth muscle cells
XPS: X-Ray photoelectron Spectroscopy

## Supplemental material

**Table SI.**
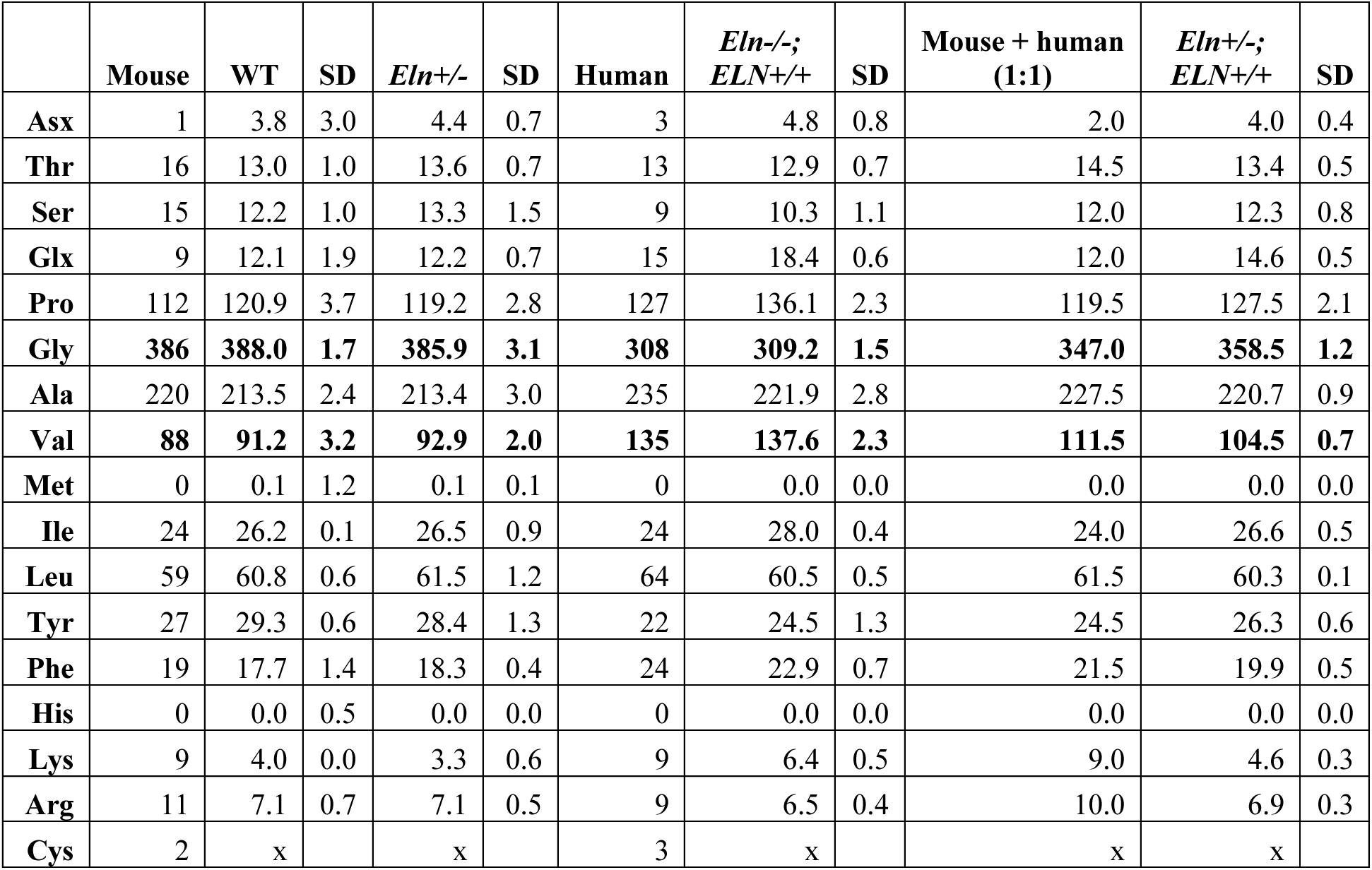
Amino acid composition of elastin in WT, *Eln+/−*, *Eln−/−;ELN+/+* aorta. The number of each amino acid/1000 residues is shown.

**Table SII.** Raman peak assignments for calcified aortas from *Mgp−/−;Eln−/−;ELN+/+* mice.

| Peak (cm<sup>-1</sup>) | Assignment |
| --- | --- |
| $\nu$ C=O (amide I) | 1658 |
| $\delta_{as}$ CH<sub>2</sub>, CH<sub>3</sub> (proteins) | 1443 |
| $\delta_s$ CH (lipids) | 1303 |
| $\delta_s$ N-H (amide III) | 1252 |
| $\nu_1$ CO<sub>3</sub><sup>2-</sup> | 1103 |
| $\nu_1$ CO<sub>3</sub><sup>2-</sup> | 1086 |
| $\nu_3$ PO<sub>4</sub><sup>3-</sup> | 1072 |
| $\nu_3$ PO<sub>4</sub><sup>3-</sup> | 1046 |
| $\nu_3$ PO<sub>4</sub><sup>3-</sup> | 1031 |
| $\nu$ C-C phenylalanine | 1003 |
| $\nu_1$ PO<sub>4</sub><sup>3-</sup> (DCPD) | 985 - 989 |
| $\nu_1$ PO<sub>4</sub><sup>3-</sup> (OCP) | 966 - 967 |
| $\nu_1$ PO<sub>4</sub><sup>3-</sup> (HA and CHA) | 959 - 962 |
| $\nu_1$ PO<sub>4</sub><sup>3-</sup> (ACP) | 941 - 953 |
| $\nu_1$ C-C (proteins) | 868 |
| $\nu_4$ PO<sub>4</sub><sup>3-</sup> | 602 |
| $\nu_4$ PO<sub>4</sub><sup>3-</sup> | 591 |
| $\nu_4$ PO<sub>4</sub><sup>3-</sup> | 561 |
| $\nu_2$ PO<sub>4</sub><sup>3-</sup> | 460 |
| $\nu_2$ PO<sub>4</sub><sup>3-</sup> | 428 |

$\nu$ : stretching; $\nu_s$ : symmetric stretching; $\delta_s$ : bending; $\delta_{as}$ : antisymmetric bending

## Notes

### Competing Interest Statement

The authors have declared no competing interest.

